# The Temporal Dynamics of Willed Attention in Vision

**DOI:** 10.1101/2022.04.11.487895

**Authors:** John G. Nadra, Jesse J. Bengson, Alexander B. Morales, George R. Mangun

## Abstract

Most models of attention distinguish between voluntary and involuntary attention, the latter being driven in a bottom-up fashion by salient sensory signals. Studies of voluntary visual-spatial attention have used informational or instructional cues, such as arrows, to induce or instruct observers to direct selective attention to relevant locations in visual space in order to detect or discriminate subsequent target stimuli. In everyday vision, however, voluntary attention is influenced by a host of factors, most of which are quite different from the laboratory paradigms that utilize attention-directing cues. These factors include priming, experience, reward, meaning, motivations, and high-level behavioral goals. Attention that is endogenously directed in the absence of external cues has been referred to as self-initiated attention, or in our prior work as “willed attention”. Such studies typically replace attention-directing cues with a “prompt” that signals the subject when to choose where they will attend in preparation for the upcoming target stimulus. We used a novel paradigm that was designed to minimize external influences (i.e., cues or prompts) as to where, as well as when, spatial attention would be shifted and focused. Participants were asked to view bilateral dynamic dot motion displays, and to shift their covert spatial attention to either the left or right visual field patch at a time of their own choosing, thus allowing the participants to control both *when* and *where* they attended on each trial. The task was to discriminate and respond to a pattern in the attended dot motion patch. Our goal was to identify patterns of neural activity in the scalp-recorded EEG that revealed when and where attention was focused. Using machine learning methods to decode attention-related EEG alpha band activity, we were able to identify the onset of voluntary (willed) shifts of visual-spatial attention, and to determine where attention was focused. This work contributes to our understanding of the neural antecedents of voluntary attention, opening the door for improved models of attentional control, and providing steps toward development of brain-computer interfaces using non-invasive electrical recordings of brain activity.

## Introduction

William James famously wrote, *“Everyone knows what attention is. It is the taking possession by the mind, in clear, and vivid form, of one out of what seems several simultaneously possible objects or trains of thought*” (James, 1890). Attention is the cognitive ability that allows humans to ignore irrelevant stimuli and hone in on the most relevant sensory inputs. Attention may be controlled by either top-down (goal directed or *voluntary*) or bottom-up (sensory or *reflexive*) influences (Bowling, Friston, & Hopfinger, 2020; Corbetta & Shulman, 2002; Jonides, 1983; Posner, 1980).

The ability to exert voluntary control over the focus of our attention is arguably a key component of the integrated sense of being that humans experience (Posner, 1994). For decades, voluntary attention has been effectively studied in humans in attention cuing paradigms using behavioral, electroencephalographic (EEG), and neuroimaging methods (Corbetta, Kincade, Ollinger, McAvoy, & Shulman, 2000; Harter, Anllovento, & Wood, 1989; Harter, Miller, Price, Lalonde, & Keyes, 1989; Hopfinger, Buonocore, & Mangun, 2000; Luck, Hillyard, Mouloua, & Hawkins, 1996; Mangun & Hillyard, 1991; Posner, Snyder, & Davidson, 1980). In such paradigms, the experimenter determines how the observer will allocate their attention by manipulating their expectancy about when, where or what an upcoming task-relevant target may be (e.g., Kingstone, 1992; H. J. Muller & Rabbitt, 1989; Posner et al., 1980), or instructing the observer how to focus attention on each trial (e.g., Hopf & Mangun, 2000; Hopfinger et al., 2000; Mangun & Buck, 1998). Arguably, however, studying voluntary attention with cuing paradigms does not capture the full range of causal influences involved in attention behavior.

In everyday vision, *voluntary* attention is biased by many factors, most of which are quite different from the highly-controlled paradigms (e.g., cuing paradigms) utilized in the laboratory. Factors biasing attention include priming (Li, Wolfe, & Chen, 2020), experience (Brockmole & Henderson, 2006; Goldfarb, Chun, & Phelps, 2016; Theeuwes, 2019; van Moorselaar, Daneshtalab, & Slagter, 2021), reward (Della Libera & Chelazzi, 2009; Failing & Theeuwes, 2018; Hickey, Chelazzi, & Theeuwes, 2010; Meyer, Sheridan, & Hopfinger, 2020; Peck, Jangraw, Suzuki, Efem, & Gottlieb, 2009), object meaning (Gayet & Peelen, 2022; Hayes & Henderson, 2021; Peacock, Cronin, Hayes, & Henderson, 2021) and high-level behavioral goals and motivations (Banerjee, Frey, Molholm, & Foxe, 2015; Lepsien, Thornton, & Nobre, 2011; Luck, Gaspelin, Folk, Remington, & Theeuwes, 2021; McMains & Kastner, 2011; Serences et al., 2005). When attention is voluntarily directed in the absence of explicit external cues, this has been referred to as internally-driven (Taylor, Rushworth, & Nobre, 2008) or self-initiated (Hopfinger, Camblin, & Parks, 2010) attention, or in our work as “willed attention” (Bengson, Kelley, Zhang, Wang, & Mangun, 2014; Bengson, Kelley, & Mangun, 2015; Bengson, Liu, Khodayari, & Mangun, 2020; Liu et al., 2017; Rajan et al., 2018). The idea is that volition drives attention in manner analogous to the volitional motor actions in studies of movement intention (Haggard, 2008; Haynes et al., 2007; Libet, Wright, & Gleason, 1983; Soon, Brass, Heinze, & Haynes, 2008), but is, arguably, theoretically dissociable (Searle, 1980). Willed attention is of particular utility when behavioral goals are in conflict with bottom-up salience and other attention-biasing influences (Bacon & Egeth, 1994; Lavie, 2005; Mevorach, Hodsoll, Allen, Shalev, & Humphreys, 2010; Theeuwes, 2018).

Although behaviorally, cued and willed attention can result in very similar patterns of reaction time and accuracy for cued/attended and uncued/ignored target stimuli (Bengson et al., 2014), cognitive neuroscience measures have revealed distinct neural signatures that characterize cued and willed spatial attention both prior to the allocation of attention (Bengson et al., 2014), and after (Bengson et al., 2015; Bengson et al., 2020; Hopfinger et al., 2010; Liu et al., 2017; Rajan et al., 2018; Taylor et al., 2008). In these willed attention studies, which are derivative of traditional spatial cuing paradigms, willed attention is typically engaged by presenting a stimulus that serves as a “prompt” that signals the subject that they should voluntarily choose where to attend on that trial (Bengson et al., 2014; Hopfinger et al., 2010; Taylor et al., 2008). Looking prior to the prompt, Bengson and colleagues were able to reveal the patterns of spontaneous brain activity (EEG alpha power) immediately before (within 1000 msec) the subjects decision about where to attend that predicted those decisions (Bengson et al., 2014). Importantly, because the prompts were lower probability (33% of trials, the other 66% being spatial cue trials), and appeared following a 2 to 8 sec jittered inter-trial-interval, the patterns of brain activity observed could not be reflections of pre-planned decisions about where to attend on each trial. Here, we continue the investigation of the antecedent brain states to willed attention in order to determine whether such patterns of brain activity can reveal <u>when</u> as well as where subjects focus spatial attention. We do this using an experimental paradigm where no cue or prompt is present.

In order to investigate the neural antecedents of willed attention, EEG was recorded from volunteers while they participated in a novel spatial attention paradigm. The participants viewed two uncued, lateralized dynamic random dot motion stimulus displays, and were instructed to choose whether and when to covertly attend the left or right display in order to discriminate features of the dynamic display, which required a button press. The goal of the study was to eliminate any cue, even a temporal one, from the task, with the aim of using the EEG measures to identify both where and when the subjects allocated their spatial attention. Isolating the neural signature(s) of self-generated, decision-driven mechanisms of spatial attention would not only further our understanding of intention, but could yield a method to enable the brain’s attention-related activity to be detected for the control of external devices, such as in brain-computer interface (BCI) applications, and therefore would be a significant contribution to both basic and applied cognitive neuroscience.

## Methods

### Participants

EEG data was recorded from 28 undergraduate student volunteers (20 female, 8 male) at the University of California, Davis. All participants had normal or corrected-to-normal vision, gave informed consent, were screened for neuropsychiatric conditions, and were paid for their participation. One subject was removed for an inability to following the task instructions, another was removed for a technical issue with data collection, four subjects were removed for excessive EEG artifacts contaminating more than 25% of their data, and two subjects were removed as they had no trials remaining in at least one bin after artifact rejection when conducting the reaction time analyses. Thus, the final analysis was conducted on 20 right-handed subjects who met all inclusion criteria.

### Paradigm and Stimuli

Each trial began with the presentation of a circular patch of 250 red and blue dots in each hemi-field (**Figure 1**). Each patch of dots had a radius of 5 degrees of visual angle, and each dot was approximately 0.23 degrees of visual angle. Each patch was located on the horizontal meridian, approximately 4 degrees (to center) lateral to fixation. In order to enable the possible analysis of focused attention using the steady state visual evoked potential (SSVEP) method (M. M. Muller, Teder-Salejarvi, & Hillyard, 1998), in one hemi-field flickered continuously at 4 Hz, while those in the other flickered at 6 Hz. From trial-to-trial, on a random basis, the frequency of flicker in the left and right patches varied (one patch always 4 Hz and the other 6 Hz); the SSVEP data is <u>not</u>, however, considered in this report. The dots varied randomly in position by 0.08 degrees every one to three screen refreshes (16.67 ms), which induced the perception of continuous random motion. In addition, within each hemi-field the proportion of red to blue dots varied in a systematic and continuous fashion from a minimum of 20 red dots in the center with 230 blue dots surrounding, to a maximum of 230 red dots in the center and 20 red dots in the surround.

**Figure 1.**
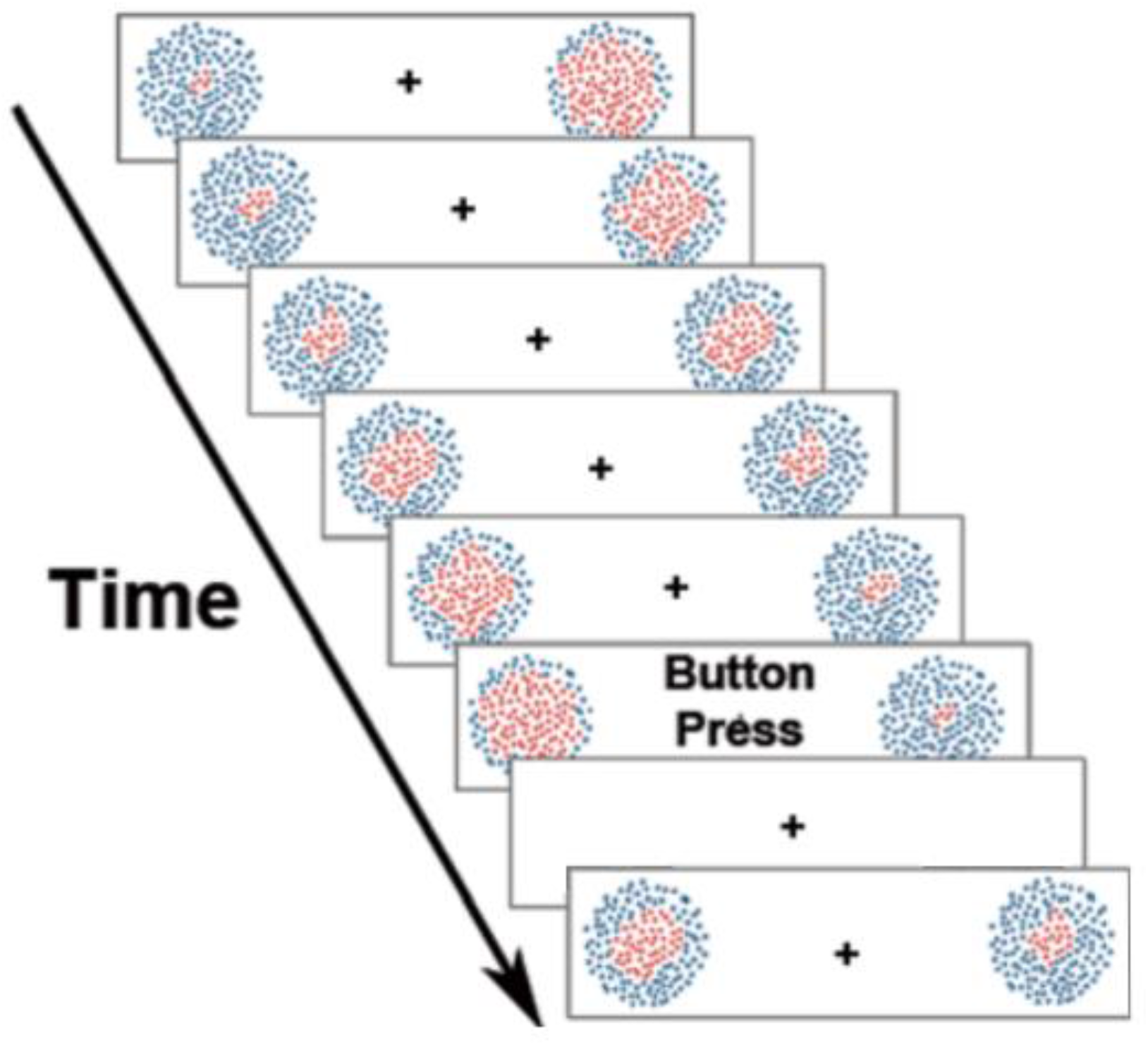
Diagrammatic representation of the dynamic stimulus arrays. An illustration of the ongoing sequence of screen refreshes, each panel illustrating one refresh. When the sequence offsets, there was a constant 500 ms delay between the offset and onset of the next trial.

With each screen refresh the number of red dots increased by 4 as a growing circle, and the number of the blue dots decreased by 4 as a decreasing annulus; this created the impression of an expanding circle of red dots within the field of blue dots in each circular patch. Once the red dot number reached the maximum of 230 red dots, the pattern changed directions so that red dots started to decline (being replaced by blue). The perception of dot patches is of a continuously expanding and contracting circle of red dots within each circle. Given a 16.67 ms refresh rate, the time from minimum to maximum in the number of red dots was approximately 1.0 sec, but, on average, each trial lasts approximately 4 seconds, because subjects could begin covert attention at varying intervals after the onset of the array. The expansion/contraction of the red dots in the left and right hemifields occurred asynchronously, so that it was not possible to predict what the pattern in one hemifield was doing given that in the other hemifield. After a button press, the patches disappeared for 500 ms, and the next trial began when the patches reappeared. The fixation point remained on the screen for the duration of each block.

### Procedure

Participants were instructed to maintain ocular fixation on the center cross and to not deviate their eyes. They were asked to voluntarily select one side of the bilateral display to attend at any point within the trial period, and to covertly attend the dynamic dot patches in order to detect target stimulus (maximal expansion of the proportion of red dots in the patch). They were urged to deploy their attention whenever they wished, and to maintain covert attention until the trial was completed. Importantly, they were told not to use any explicit strategy or develop any pattern for choosing when or which side to deploy covert attention (such as alternating sides on each trial), and to not decide prior to trial onset which hemifield patch to attend. In other words, once the bilateral array appeared, the subjects were requested to make a spontaneous decision during each trial about which side to focus covert spatial attention. The participants were told to maintain their attention on the chosen hemi-field patch for at least one full expansion cycle (approximately one second) while trying to discriminate the maximum size of the expanding red dots in the chosen hemi-field. The participants were required to press a button (either the left arrow or right arrow on a keyboard) as quickly as possible when they perceived that the red dots were at their maximal expansion in the attended hemifield only, and were told to completely ignore the opposite hemifield patch.

### EEG Recording and Analysis

The EEG was recorded from 64 tin electrodes (mounted in an elastic electrode cap; Electro-cap Int.) at the following scalp locations: Fp1, Fp2, F7, F3, Fz, F4, F8, FC5, FC1, FC2, FC6, T7, C3, Cz, C4, T8, CP5, CP1, CP2, CP6, P7, P3, Pz, P4, P8, PO9, O1, Oz, O2, PO10, AF7, AF3, AF4, AF8, F5, F1, F2, F6, FT9, FT7, FC3, FC4, FT8, FT10, C5, C1, C2, C6, TP7, CP3, CPz, CP4, TP8, P5, P1, P2, P6, PO7, PO3, POz, PO4 and PO8 (Oostenveld & Praamstra, 2001). These sites were referenced to FCZ during recording, but were re-referenced offline to the algebraic average of TP9 and TP10 (adjacent to the left and right mastoids). The continuous EEG was recorded with a bandpass of DC-100 Hz and digitized at 1000 samples per second, and then downsampled offline to 250 samples per second. Before artifact rejection, a bandpass filter for 0.05 to 50 Hz was applied to the data. Eye-blinks were removed using independent component analysis (ICA) methods (Vigario, 1997). Residual artifacts were detected automatically, and trials with excessive artifacts were removed using ERPLAB’s moving window peak-to-peak artifact rejection (100 μV threshold), iterating through the data with a moving window of 100 ms in 50 ms steps. An additional moving window approach was applied to channels FT9 and FT10 to ensure no trials with eye movements were left in the data. The parameters for this additional moving window approach marked all trials which exceeded 20 μV within a sliding 100 ms window, across 50 ms steps. Each epoch was also visually inspected to manually reject artifacts not picked up by the prior methods, as well as verify that the artifact rejection pipeline was functioning as intended. The data was then epoched in two separate time periods, -1000 ms to 4000 ms relative to the onset of the sequence, as well as -4000 ms to 1400 ms relative to the button press. In the event-related potential analysis, a baseline from -500 ms to 0 ms relative to the onset of the flickering patches (trial onset). For the stimulus onset data, only trials where participants reported their shift of attention within 4,000 ms were retained. Three subjects did not have any data remaining within this range, so their data was excluded from further analysis. Pre-processing was conducted using both the EEGLAB (Delorme & Makeig, 2004) and ERPLAB (Lopez-Calderon & Luck, 2014) plugins for MATLAB.

In line with prior work on willed attention (Bengson et al., 2014), we focused our analyses on the alpha band of the EEG. To examine the onset and strength of the willed attention signal, we extracted the trial-by-trial alpha band signal relative to two distinct time points; (i) the onset of the sequence and moving forward in time, as well as (ii) moving backwards in time from the onset of the button press, which logically followed willed shifts of covert attention. We then used the direction of the decision to attend as a grouping variable, labelling trials relative to whether the participant chose to attend to the left or right hemifield on a trial-by-trial basis. For these analyses, the time-frequency analysis was performed on each trial via a short sliding Hanning taper with an adaptive time window of three cycles at each frequency - conducted from 9-11 Hz. The alpha frequency band analysis was placed at 9-11 Hz to minimize overlap with the 4 Hz and 6 Hz background flickering of the stimulus arrays. The analysis was conducted using the fieldtrip toolbox plugin for MATLAB (Oostenveld, Fries, Maris, & Schoffelen, 2011).

We implemented a support vector machine (SVM) decoding pipeline that was similar to that utilized by Bae and Luck (Bae & Luck, 2018). The fitcsvm() function in MATLAB was used to carry out this analysis. A 3-fold cross validated support vector machine was trained and tested separately over each individual time point (in 20 ms increments). The cross-validation that was implemented allowed the same data to act as both the training and testing sets. The data was split into three equal portions, where in the first iteration, two thirds are used for training, then one third is used for testing. On the next iteration, the training and testing sets are randomized, to test the classifier across varying subsets of the data. This process was repeated across ten iterations, then the accuracies obtained from the testing set were averaged across iterations for each time point. The data was averaged over trials, 19 relevant electrode channels were included (all parietal and occipital electrodes) and the data was fourier transformed (extracting alpha band signals at 9-11 Hz) before the training and testing of the SVM. The classification was a binary SVM, computing the classification accuracy of trials where subjects were deploying attention to the left versus the right. Given trial count (i.e., attending left vs. right) is potentially variable given self-generated decisions about where to attend, each subject’s trial count per bin (left/right) was set to equal trial lengths by removing the last trials of the larger bin. We used a nonparametric cluster-based Monte Carlo simulation technique (similar to the commonly used cluster-based mass univariate approach). This method was chosen due to its correction for multiple comparisons and the fact that decoding accuracy may not be normally distributed. The decoding accuracy was extracted at each timepoint, then tested with a one-sample t-test (one sample, as below chance decoding is not relevant to our findings). We then searched for significant clusters where the t-tests were significant (p < 0.05) and the t scores were combined to create a cluster-level t score. Then we assessed whether the cluster t score was higher than the t score expected by chance (generated by the Monte Carlo simulation), which controls the type 1 error rate at a cluster level. Then, each simulated trial was a randomly sampled number (whether one or two) to compute the chance level for each bin (left or right). The Monte Carlo technique had 10 iterations, with three validations, indicating this process was repeated 60 times (2 bins x 3 cross validations x 10 iterations). This process was then repeated once for each time point (201 time points for the sequence onset decoding; 176 for the button press decoding), to find an accurate decoding accuracy at each datapoint. The data was then smoothed over five timepoints for graphing purposes. This process was repeated for each of our subject’s data.

## Results

### Reaction Time

The mean reaction time (RT)—as measured from trial onset—was 3,945 ms for trials where attention was deployed to the left, and 3,908 ms when to the right. The difference between these reaction times was not statistically significant in a two-sample t-test (p = 0.6289). Given the task design (unattended-sided stimuli were to be completely ignored), there are no behavioral measures of selective spatial attention (i.e., attended vs. unattended RTs); however, see the alpha band topographic analysis below, which addresses the issue of whether subjects were focusing spatially selective attention in this task. In terms of how many times participants chose to attend to each side, participants reported (via their left or right button press) that they chose to covertly attend the left hemifield patch 4,924 times in total, while the right hemifield patch was attended 4,653 times across all participants. To examine whether the previous trial influenced decision outcomes, a logistic regression generalized linear model was fit to the behavioral data, which solidified that the previous trial did not have an influence on the direction that attention was shifted on any given trial (p = 0.3375).

### Alpha-Band Oscillations

We began by validating our task in order to ensure that subjects had allocated selective visual-spatial attention in our design. To do this we relied on the well-established EEG alpha correlates of focused visual spatial attention which show left versus right posterior scalp EEG alpha power asymmetries with spatial attention to lateral visual field locations (Bengson et al., 2014; Liu, Bengson, Huang, Mangun, & Ding, 2016; Rihs, Michel, & Thut, 2007; Romei, Gross, & Thut, 2010; Worden, Foxe, Wang, & Simpson, 2000). We compared the distribution of alpha power across the left and right posterior scalp for the choose-left and choose-right trials (Fig. 3). We found significant (two-sided t-test) left versus right alpha power asymmetries over posterior scalp in the 1000 ms prior to the button press (p < 0.01); this pattern was not significant in an earlier time window from -2000 to -1000 ms prior to the button press (p = 0.1116). This alpha power lateralization with spatial selective attention demonstrated that alpha-band oscillations serve as a reliable index of the direction of covert spatial attention in our willed attention design. With this expected result firmly established, we turned to decoding the time course of the allocation of willed attention.

**Figure 2.**
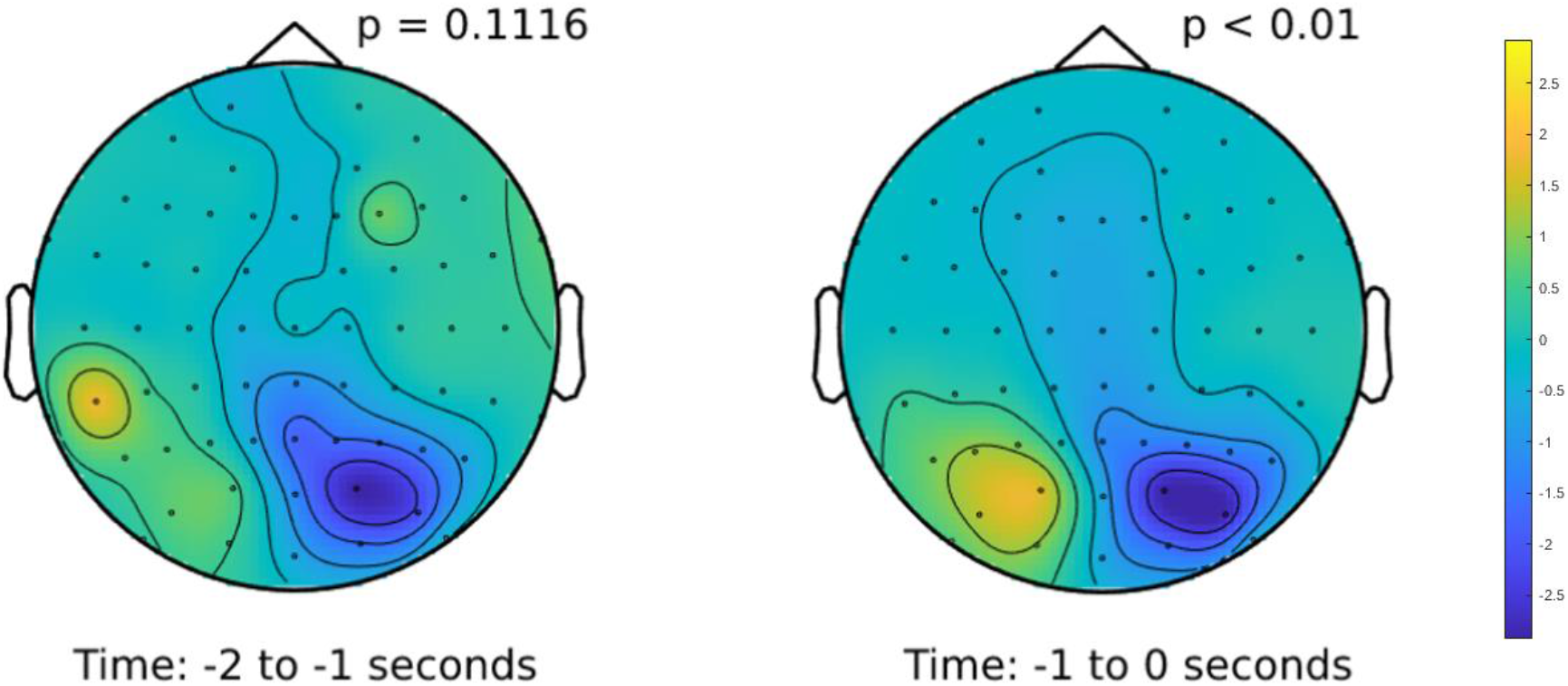
Alpha-band (9-11Hz) difference plot (attend left - attend right) during two distinct time periods preceding the button press to mark the active focus of attention to the chosen hemifield. The left panel shows the time period from -2 sec to -1 second before the button press, while the right panel shows the time period directly preceding the report, from one second before the button press to the onset of the button press itself. The color scale is based on the absolute value relative to the highest/lowest difference in power across both plots.

**Figure 3.**
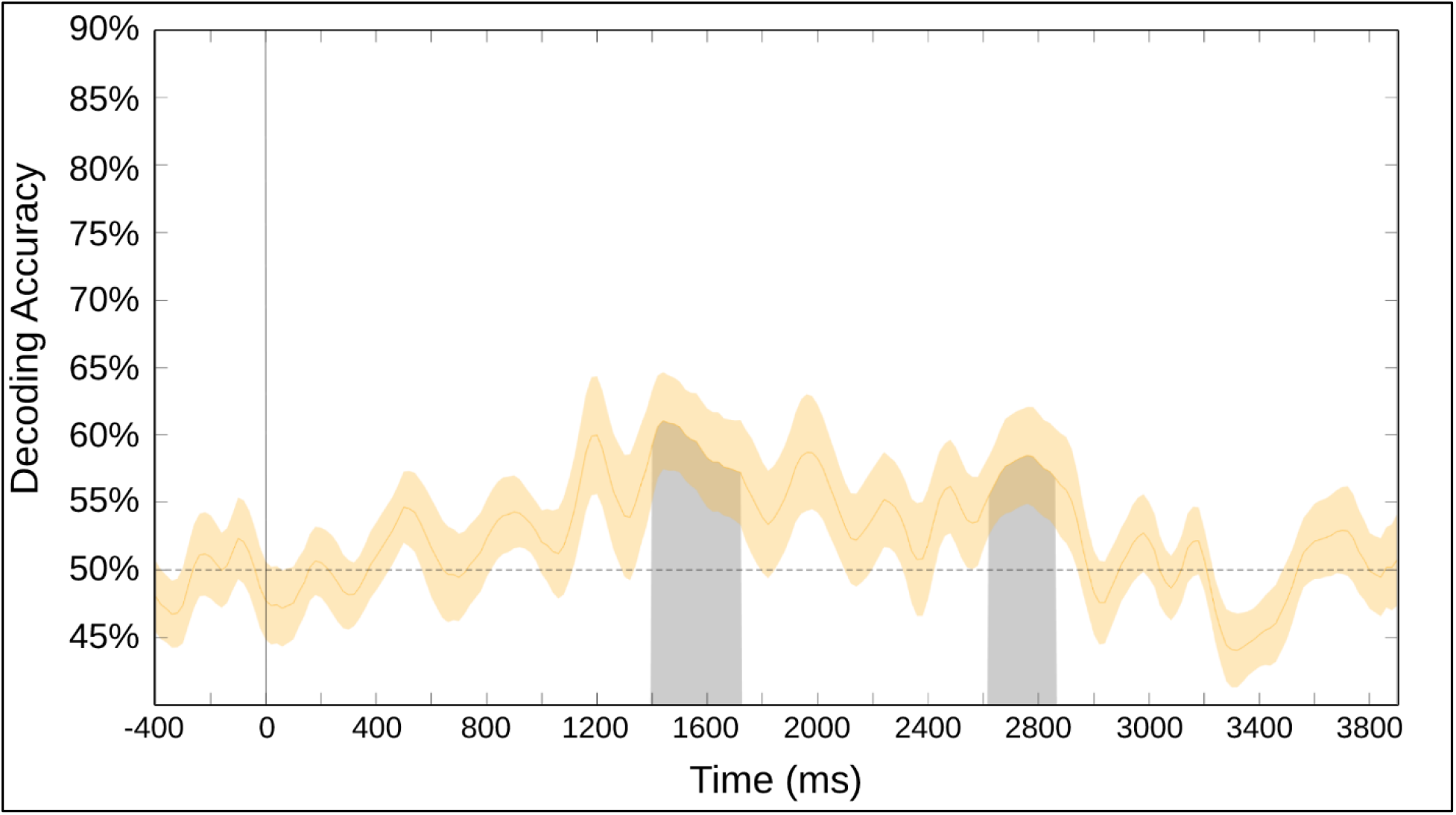
This data is epoched relative to the sequence onset (represented at time 0 by a vertical line). Only short trials (between 500 ms and 4,000 ms) were included in the analyses, which is why we see a drop off around the three second mark. This analysis took place in 19 parietal/occipital electrodes.

### Decoding Results

Figure 3 shows the decoding accuracy for attend-left versus attend-right during willed attention, collapsed across the 17 subjects in the study. These SVM classifier results are for the data epoched to the onset of the bilateral array (t=0 ms). Decoding accuracy starts at chance level (dashed line) and gains rising about chance over time. The decoding accuracy rose above chance starting at∼1,400 ms after the array onset and lasted until∼1,700 ms. Following a dip in decoding accuracy, a long-latency period also shows statistically significant decoding accuracy (∼2,625-2,850 ms after array onset). This conceptually lines up with our expectations, as we have only included trials with short reaction times, where participants have reported their decision to attend within 4,000 ms after the onset of the array.

The support vector machine classifier results for data epoched to the button press show significant decoding throughout the entire period from 1,900 ms before, spanning∼750 ms after the button press (fig. 4). There are also patches of significant decoding, as also similar to the results shown in the decoding period relative to the sequence onset (Fig. 3). We then see a gradual ramp down to chance as time progresses (∼750 ms post-button press). These results accurately reflect the goal of the experimental design where participants have the freedom to choose how long they are attending to each side freely and serves as an example of the reliability of the alpha-band fluctuations present in fully volitional decisions to attend. With these results, we have demonstrated the capability to achieve significantly above average decoding of

**Figure 4.**
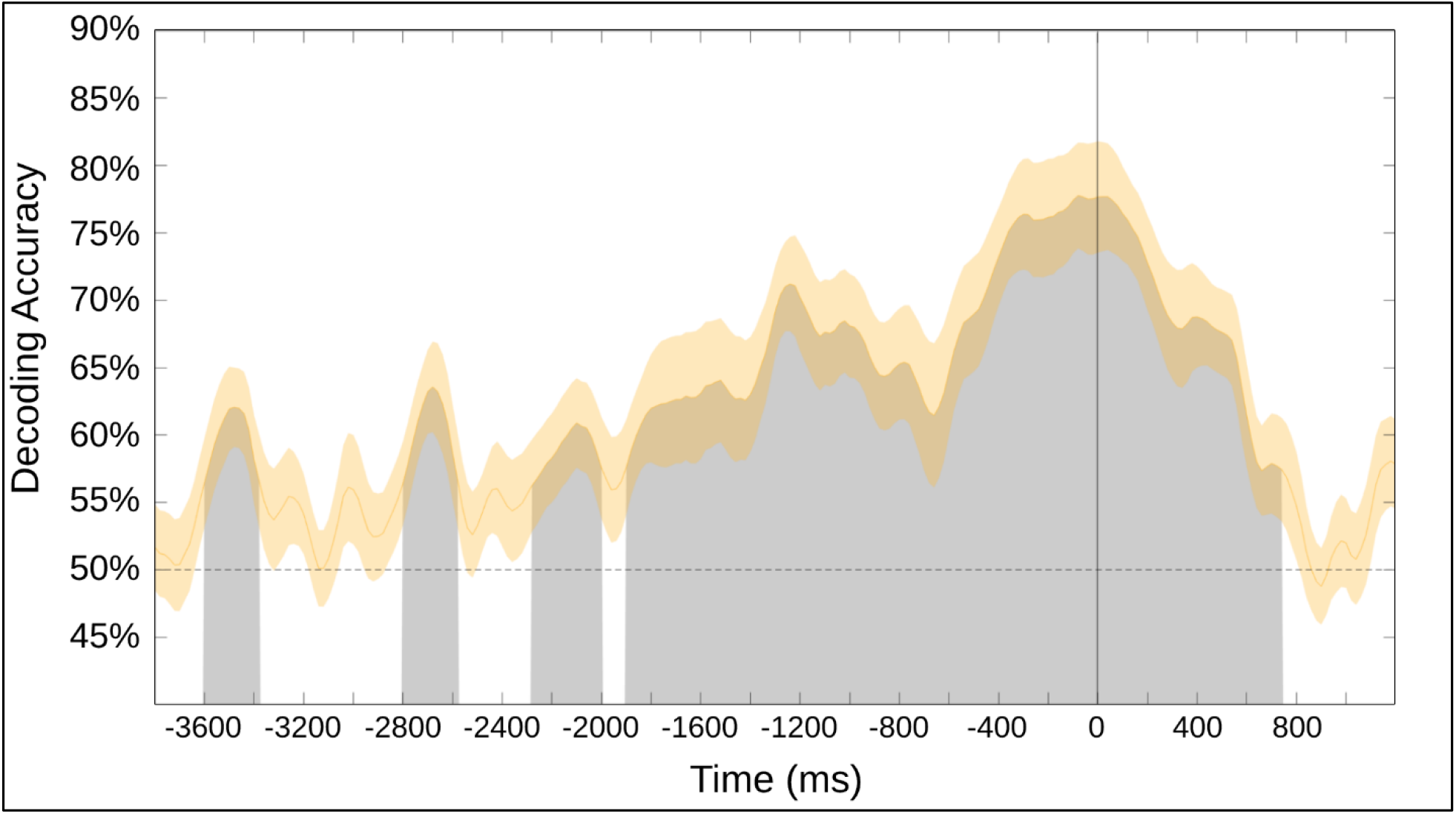
The support vector machine decoding accuracy at each time point, epoched relative to the button press signaling sustained attention to one hemifield. This analysis was done over 19 occipital electrodes. Black line denotes time zero, which is the recorded onset of the button press.

sustained volitional attention in a setting with minimal external influence or cueing, as well as the ability to decode the average onset of a volitional shift in spatial attention.

## Discussion

In this study, we investigated voluntary visual-spatial attention when guided internally by the subject’s choices about both *when* and *where* to focus attention (i.e., willed attention). Prior research on willed attention asked subject to choose where to attend, but always in response to a prompt that signaled the subjects <u>when</u> to voluntarily engage spatial attention (Bengson, Kelley, & Mangun, 2015; Bengson et al., 2014; Bengson et al., 2020; Hopfinger et al., 2010; Liu et al., 2017; Rajan et al., 2018; Taylor et al., 2008). The act of deploying attention in the real world need not be cued externally; that is, not every shift of attention requires an explicit extrinsic temporal or spatial guiding signal (Bengson et al., 2014; Hopfinger et al., 2010; Taylor et al., 2008). In the present work, we presented subjects with bilateral, dynamic dot motion displays, asking them to view the displays, and then to spontaneously focus spatial attention on either the left or right patch at a time of their choosing, thus eliminating any explicit external attentional cue or prompt.

We focused our analyses on EEG alpha oscillations (9-11 Hz), which has been implicated in many forms of attention (for a review, see, Van Diepen, Foxe, & Mazaheri, 2019). It is well known that the selective deployment of spatial attention in the lateral visual fields in response to attention-directing cues is correlated with lateralized changes in EEG alpha power over the occipital scalp (Liu et al., 2016; Rihs et al., 2007; Thut, Nietzel, Brandt, & Pascual-Leone, 2006; Voytek et al., 2017; Worden et al., 2000). In the present study, we used decoding of EEG alpha patterns to investigate the time course and locus of willed attention shifts within uncued, dynamic visual displays. Decoding has proven useful for understanding the contributions of EEG alpha signals to spatial attention in prior research (Bae & Luck, 2018; Samaha, Sprague, & Postle, 2016).

We found that volitional attention can be actively decoded from strictly EEG data in the alpha-band range (9-11Hz). By epoching our analyses relative to the onset of a trial, we have isolated the electrophysiological activity related to a self-generated shift in covert attention, thus building upon former studies on willed attention (Bengson et al., 2014). We also epoched our data relative to a button press, signaling covert attention has been shifted and active attention is deployed to a specific hemifield. This analysis has shown multiple significant bumps in decoding accuracy between 2-4 seconds before the button press, which is followed by a larger time span of significance related to the active deployment of sustained attention to one hemifield, remaining about 750 milliseconds after the button press before tapering back to chance-level decoding accuracy. The evidence provided here establishes the ability to pinpoint the average onset of a shift in volitional attention, as well as highlights the importance of the willed attention signal in exhibiting the volitional control of attention, even without the constraints of a typical cue-to-target paradigm.

Self-generated processes have primarily been studied in the context of motor intention and actions (for a review, see, Eagleman, 2004). In this area of scholarship, a distinction has been drawn between willed and automatic control of actions, with attention being a key distinguishing component of prominent models (Norman & Shallice, 1986; Shiffrin & Schneider, 1977). A core concept is that intentions arise prior to actions, and that the antecedent neural activity could therefore provide information about the underlying neural mechanisms of intention. For example, Benjamin Libet’s work on motor intentions sought to reveal the neural correlates of intentions to act (Frith & Haggard, 2018; Libet, Gleason, Wright, & Pearl, 1983; Libet, Wright, & Gleason, 1982). His subjects were instructed to make a volitional muscle movement at a time of their choosing while watching a pseudo clock face on an oscilloscope screen on which a dot rapidly swept around the circular screen in clockwise fashion. They were told to report the “time” on the pseudo clock face at which they first were aware of their intention to act. Libet used the reported value as a time stamp that he compared to the backwards averaged event-related potentials (ERP) that were time-locked to the motor action (indicated by the onset of electromyographic activity of the relevant forearm muscle). He found that the antecedent ERP activity preceded the reported time of first intention by hundreds of milliseconds. Our present study applied a similar framework as this literature on self-generated motor actions, but instead probed willed attention by backward decoding the EEG from the button press response, allowing us to establish the onset of willed attention as the time period in which the decoding accuracy for left versus right choices rose above chance.

Our findings have direct consequence for our understanding intention by moving beyond the oft-studied realm of intentions to make a movement, to the case of intention to attend, that is, willed attention. Importantly, it demonstrates that even in cases where the subject is not prompted to make a decision, fully self-paced decisions have decodable neural correlates. In our work, based on scalp-recoded EEG, we are unable to identify the underlying functional anatomical correlates of the decision about when and where to attend, but our approach may nonetheless provide directions for future research, for example, using magnetoencephalography (MEG) (Hardy, Jensen, Wheeldon, Mazaheri, & Segaert, 2022) or intracranial recording (Helfrich et al., 2018; Stolk et al., 2018)

This general approach may also be generally applicable in applied research, for example, in BCI applications, where brain activity related to intentions to attend could be tapped to control devices by inferring intentions directly. A BCI should be built around neural signals having reliable features for feature extraction (i.e., they should reflect the subjects’ intent), would benefit if based on a non-invasive technique (e.g., scalp-recorded EEG), and should also have optimal signal-to-noise (Choi & Kim, 2019; O’Sullivan et al., 2015; Shih, Krusienski, & Wolpaw, 2012). Alpha oscillations elicited by a decision to attend in a willed attention setting may be such a signal, being recordable non-invasively from the scalp (as well as intracranially), and having a relatively high signal-to-noise (i.e., alpha-to-ongoing EEG) ratio. Establishing the reliability of the alpha signal as a measure of intention without the constraint of external cuing or prompting, is a positive step forward in this regard. Previous research using EEG has established the ability to decode binary covert intention on a single-trial basis (Choi & Kim, 2019), and applying such methods to willed attention data such may provide the foundation for future BCI applications that tap intentions in cognitive acts.

## Acknowledgements

This work was supported by MH117991 to George R. Mangun and Mingzhou Ding, and NSF BCS 1339049 to George R. Mangun and Jesse J. Bengson. We are grateful to Sharon Corina, Ali Mazaheri and Carlos Carrasco for their support and advice, as well as Manvita Tatavarthy for their assistance with data collection.

## Notes

**Conflict of Interest:** The authors declare no competing financial interests.

### Competing Interest Statement

The authors have declared no competing interest.

